# HSI2/VAL1 and HSL1/VAL2 function redundantly to regulate seed dormancy by controlling *DOG1* expression in Arabidopsis

**DOI:** 10.1101/2019.12.20.885392

**Authors:** Naichong Chen, Hui Wang, Haggag Abdelmageed, Vijaykumar Veerappan, Million Tadege, Randy D. Allen

## Abstract

*DELAY OF GERMINATION1* (*DOG1*) represents a major quantitative locus for the genetic regulation of seed dormancy in Arabidopsis. Accumulation of DOG1 in seeds leads to deep dormancy and delayed germination. Here, we report that the conserved B3 DNA binding domains of the transcriptional repressors HIGH-LEVEL EXPRESSION OF SUGAR INDICIBLE GENE2/ VIVIPAROUS-1/ABSCISIC ACID INSENSITIVE 3-LIKE1 (HSI2/VAL1) and HSI2-LIKE1/ VIVIPAROUS-1/ABSCISIC ACID INSENSITIVE 3-LIKE2 (HSL1/VAL2), which play critical roles in the developmental transition from seed maturation to seedling growth, interact with RY elements in the *DOG1* proximal promoter leading to repression of *DOG1* transcription during germination and seedling establishment. *DOG1* expression is partially de-repressed in *hsi2/val1* (*hsi2-2*) but not in *hsl1/val2* (*hsl1-1*) knockout mutants and is strongly upregulated in a *hsi2/val1 hsl1/val2* double mutant, indicating that HSI2/VAL1 and HSL1/VAL2 act redundantly to repress *DOG1* expression. HSI2/VAL1 and HSL1/VAL2 form homo- and hetero-dimers *in vivo*, and dimerization is dependent on the HSI2/VAL1 PHD-like domain. Complementation of *hsi2-2* with HSI2/VAL1 harboring a disrupted plant homeodomain (PHD)-like domain results in stronger de-repression of *DOG1* expression than the *hsi2-2* knockout, indicating that the PHD-like domain plays a critical role in mediating functional interactions between HSI2/VAL1 and HSL1/VAL2. Both HSI2/VAL1 and HSL1/VAL2 interact with components of polycomb repressive complex 2 (PRC2), including CURLY LEAF and MULTICOPY SUPPRESSOR OF IRA1 (MSI1), along with LIKE HETERCHROMATIN PROTEIN 1 (LHP1), which are involved in the deposition and expansion of histone H3 lysine 27 trimethylation (H3K27me3) marks in repressive chromatin. Thus, HSI2/VAL1 HSL1/VAL2-dependent recruitment of PRC2 leads to silencing of *DOG1* through the deposition of H3K27me3.

## Introduction

Seed dormancy is an adaptive mechanism that allows for the dispersal and survival of seeds over distance and time, and ensures that germination occurs under favorable conditions (Finch-Savage et al., 2006). Seed dormancy is a complex trait that is regulated by both phytohormones and genetic factors. Abscisic acid (ABA) plays an important role in initiating and enhancing seed dormancy, while the dormant state is reversed by gibberellins, which promote germination under favorable conditions (Koornneef et al., 2002; Liu et al., 2010). Recently, *DELAY OF GERMINATION1 (DOG1)* was reported to be a major quantitative trait locus for the genetic regulation of seed dormancy in Arabidopsis (Alonso-Blanco et al., 2003; Bentsink et al., 2006, 2010; Huang et al., 2010). Compared to wild type, loss of function *dog1-3* Arabidopsis mutant seeds germinated early, whereas, gain-of-function *dog1-5* mutant seeds showed delayed germination (Cyrek et al., 2016; Huo et al., 2016). DOG1 acts in parallel with ABA to delay germination (Graeber et al., 2014) and, since DOG1 requires the clade A PP2C phosphatases PP2C to control seed dormancy (Née et al., 2017), the ABA and DOG1 pathways converge at that level. In addition, ETHYLENE RESPONSE FACTOR12 (ERF12), and its downstream target ETHYLENE RESPONSE1 (ETR1), directly bind the *DOG1* promoter and recruit TOPLESS (TPL), leading to repression of *DOG1*, suggesting that ethylene also regulates seed dormancy through the ETR1-ERF12/TPL-DOG1 module (Li et al., 2019). Low temperature during seed maturation increases *DOG1* expression and induces seed dormancy (Chiang et al., 2011; Kendall et al., 2011; Nakabayashi et al., 2012). Seed development under low-temperature conditions triggers *DOG1* expression by increasing the expression and protein abundance of bZIP67, which directly targets the *DOG1* promoter and activates *DOG1* expression, leading to enhanced seed dormancy (Bryant et al., 2019). *DOG1* is also regulated by other complex mechanisms that include histone modifications, alternative polyadenylation, alternative splicing, and a *cis*-acting antisense noncoding transcript (*asDOG1)* (Bentsink et al., 2006; Cyrek et al., 2016; Müller et al., 2012; Molitor et al., 2014; Fedak et al., 2016).

Two closely related proteins, HIGH-LEVEL EXPRESSION OF SUGAR INDUCIBLE2 (HSI2) and HSI2-LIKE1 (HSL1), play critical roles in the developmental transition from seed maturation to germination and seedling development and from vegetative growth to reproductive development (Tsukagoshi et al., 2005, 2007; Suzuki et al., 2007; Veerappan et al., 2012; Chhun et al., 2016; Qüesta et al., 2016; Yuan et al., 2016; Chen et al., 2018). These proteins, also known as VIVIPAROUS-1 /ABSCISIC ACID INSENSITIVE 3-LIKE1 (VAL1) and VIVIPAROUS-1 /ABSCISIC ACID INSENSITIVE 3-LIKE2 (VAL2), contain four conserved putative functional domains, including a plant homeodomain (PHD)-like, a plant specific B3 DNA binding domain, a cysteine and tryptophan-rich-zinc finger domain (CW), and an ethylene-responsive element binding factor-associated amphiphilic repression (EAR) domain. The HSI2 B3 domain binds to RY/Sph (RY) elements and is required for HSI2-dependent repression of *FLOWERING LOCUS C (FLC)* and *AGAMOUS-LIKE15 (AGL15)* (Qüesta et al., 2016; Yuan et al., 2016; Chen et al., 2018). The HSI2 PHD domain is also required for HSI2 accumulation at the *AGL15* locus and repression of *AGL15* expression (Chen et al., 2018). HSI2 and HSL1 can form homodimers and heterodimer *in vivo* and HSL1 shows partial redundancy to HSI2 in the regulation of *FLC* (Chhun et al., 2016; Yuan et al., 2016). HSI2 is reported to recruit polycomb repressive complex 2 (PRC2) to the *AGL15* and *FLC* loci (Qüesta et al., 2016; Yuan et al., 2016; Chen et al., 2018). HSI2 was also reported to bind HISTONE DEACETYLASE6 (HDA6) and MEDIATOR13 (MED13) and HSL1 was reported to interact with HDA19, indicating that these factors may also be involved in the repression of seed maturation genes during germination in Arabidopsis (Chhun et al., 2016; Zhou et al., 2013). The molecular mechanisms of HSL1 function and its relationship with HSI2 has not been fully characterized.

LIKE HETEROCHROMATIN PROTEIN1 (LHP1) recognizes H3K27me3 (trimethylation of lysine at histone H3) repressive marks through its chromodomain (Turck et al., 2007; Zhang et al., 2007; Exner et al., 2009), and interacts with MULTICOPY SUPPRESSOR OF IRA1 (MSI1) to positively recruit PRC2 to chromatin for the deposition of H3K27me3 (Derkacheva et al., 2013). LHP1 also directly interacts with RING-RAWUL proteins and EMBRYONIC FLOWER1 (EMF1, a component of PRC1), suggesting LHP1 can be present in several PRC1-like complexes and may act as a bridge between PRC1 and PRC2 (Xu et al., 2008; Bratzel et al., 2010). In Arabidopsis, LHP1 co-localizes with H3K27me3 across the genome, and is responsible for the expansion of H3K27me3 associated with the stabilization of transcriptional repression (Turck et al., 2007; Zhang et al., 2007; Exner et al., 2009). Expression of many tissue specific genes is upregulated in *lhp1* loss-of-function mutants (Lafos et al., 2011; Libault et al., 2005), with decreased levels of H3K27me3 seen at direct target gene loci, including *FLC* (Yuan et al., 2016; Veluchamy et al., 2016), indicating that LHP1-dependent gene repression correlates with deposition of H3K27me3. Molitor et al. (2014) reported that *DOG1* is negatively regulated by ALFIN1-like proteins, which contain PHD domains and can form protein complexes with LHP1 at the *DOG1* locus to replace H3K4me3 (trimethylation of lysine 4 at histone H3) marks associated with gene activation with H3K27me3, leading to its transcriptional downregulation and promotion of seed germination.

Here we report that the proximal region of the *DOG1* promoter is directly targeted by both HSI2 and HSL1. These transcriptional repressors recruit LHP1 and CURLY LEAF (CLF), and promote the deposition of H3K27me3 marks, leading to repression of *DOG1* and the early release of seed dormancy.

## Results

### HSI2 and HSL1 redundantly repress *DOG1* to regulate seed dormancy

Reverse transcription quantitative PCR (RT-qPCR) analysis showed that *DOG1* expression was not significantly affected, relative to wild-type (WT), in freshly harvested *hsl1-1* Arabidopsis seeds but its expression increased by approximately 3-fold in *hsi2-2* and by 14-fold in the *hsi2 hsl1* double mutant (Figure 1A), indicating HSI2 and HSL1 play overlapping roles in the repression of *DOG1* expression. Analysis of the rates of germination of freshly-harvested seeds from these Arabidopsis lines confirmed that loss-of-function mutations in both *hsi2* and *hsl1* are necessary for significant increases in seed dormancy to be detected (Figure 1B). The gain-of-function mutant *dog1-5*, which shows strongly delayed germination, and the loss-of-function mutant *dog1-3*, which shows early germination, served as controls in this assay. These results indicate that HSI2 and HSL1 act redundantly to repress *DOG1* expression in seeds. Though their functions overlap, the native *HSI2* gene is able to fully complement *HSL1*, while *HSL1* cannot fully replace the function of *HSI2*. Furthermore, while seed dormancy is promoted by *DOG1* expression, a relatively large increase in *DOG1* expression, as seen in the *hsi2 hsl1* double mutant, appears to be necessary to significantly affect seed dormancy.

**Figure 1.**
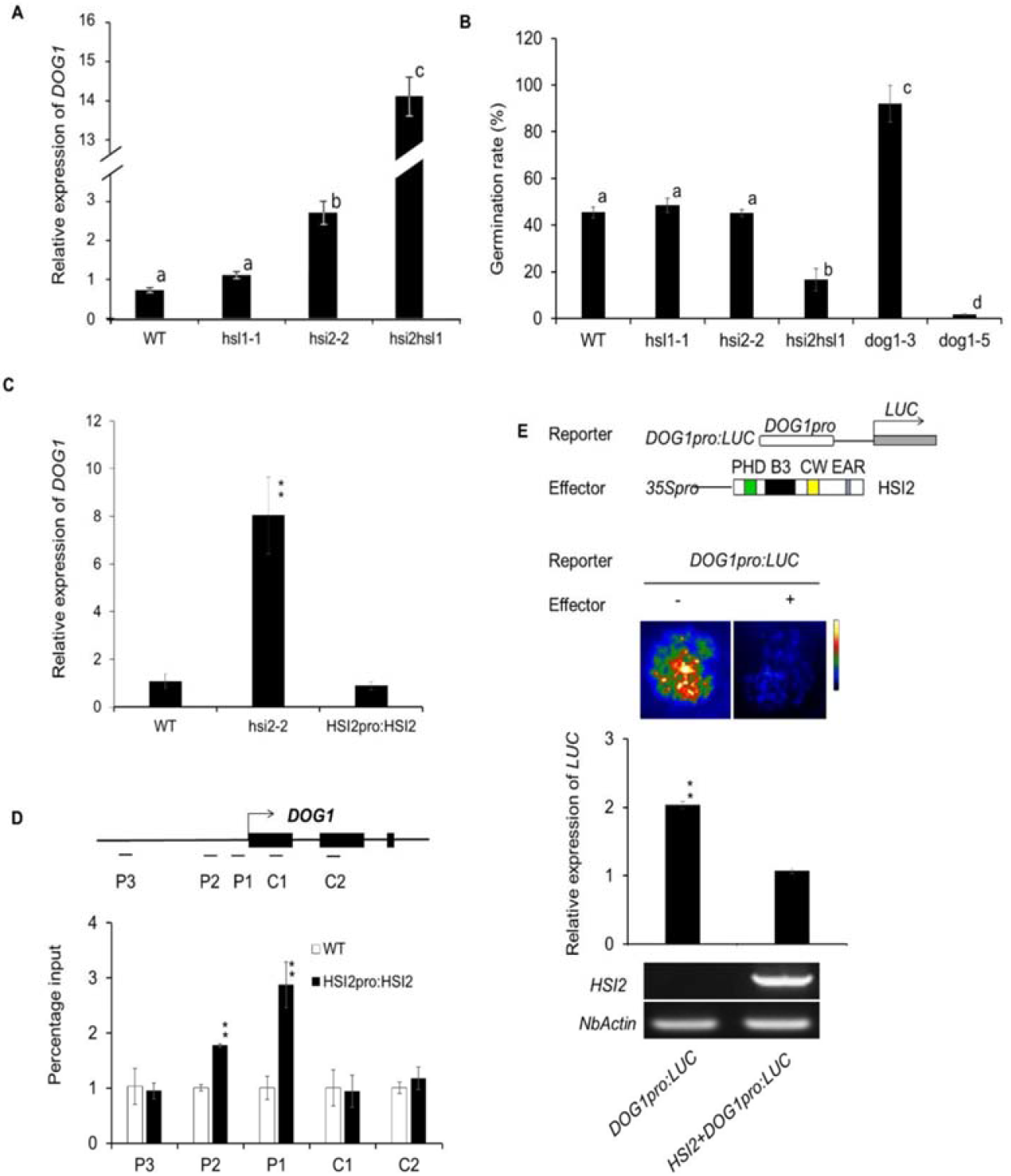
*hsi2 hsl1* double mutant shows increased primary seed dormancy and HSI2 directly binds to *DOG1* to regulate its expression. **(A)** Relative expression of *DOG1* in freshly-harvested seeds from WT, *hsl1-1, hsi2-2, hsi2 hsl1* Arabidopsis plants. RT-qPCR assays were normalized using *EF1a*. **(B)** Quantification of germination rates of freshly-harvested seeds of *hsl1-1, hsi2-2*, and *hsi2 hsl1*. Seeds from *dog1-3* loss-of-function and *dog1-5* gain-of-function mutants were used as controls. Rates of germination were scored three days after plating on half-strength MS medium. Means of three independent experiments (±SD) are shown. Lowercase letters indicate significant differences (P<0.01) between genotypes. **(C)** Relative expression of *DOG1* gene in WT, *HSI2pro:HSI2* and *hsi2-2* Arabidopsis seedlings, assayed by RT-qPCR. **(D)** ChIP-qPCR analysis of HSI2 enrichment at *DOG1* gene in WT and *HSI2pro:HSI2* Arabidopsis plants. Schematic representations of *DOG1* gene and assayed genomic regions are indicated, with P indicating promoter regions and C indicating coding regions. These regions were analyzed by ChIP-qPCR analysis of seven day old transgenic plants harboring *HSI2pro:HSI2-HA* using HA antibody. qPCR data was normalized using *ACT2* as an internal standard. Data represent means of three qPCR reactions from each of three independent ChIP assays. **(E)** Schematic representation of reporter and effector used to assay the function of the HSI2 in the regulation of *DOG1*. The reporter construct includes a luciferase gene is driven by the *DOG1* promoter. In the effector construct, expression of HSI2 is controlled by the *CaMV 35S* promoter. Luminescence images and relative expression of luciferase mRNA, determined by RT-qPCR assays, in *N. benthamiana* leaves co-infiltrated with the reporter and effector gene combinations as indicated. Expression of the *NbActin* gene was used for normalization. Error bars in indicate SD for three independent assays. ** indicate statistically significant differences (*P*<0.01) determined by Student’s *t* test.

### HSI2 directly regulates *DOG1* by binding to its promoter

Since expression of *DOG1* is upregulated and H3K27me3 enrichment at the *DOG1* locus is reduced in *hsi2-2* knockout mutant Arabidopsis seedlings (Veerappan et al., 2012, 2014), we predicted that *DOG1* could be a direct regulatory target of HSI2. To test this hypothesis, a rescued *hsi2-2* Arabidopsis line that expresses HA epitope-tagged HSI2 under the control of the native *HSI2* promoter (*HSI2pro:HSI2-HA*) was developed. Repression of *DOG1* was restored in this line with expression reduced, relative to *hsi2-2*, to levels similar to WT plants (Figure 1C). To detect and quantify the enrichment of HSI2 at the *DOG1* locus, a number of different primer sets were used to analyze promoter (P) and coding (C) regions of the *DOG1* gene by chromatin immunoprecipitation quantitative PCR (ChIP-qPCR) (Figure 1D). Significant HSI2 enrichment was detected at the proximal promoter regions (P1 and P2) of *DOG1* with the highest level of enrichment at P1. No significant enrichment of HSI2 was observed at the distal promoter (P3) or coding regions in the first and second exons (C1 and C2) of *DOG1*. To determine if transcription of *DOG1* is directly down-regulated by HSI2, a luciferase reporter gene controlled by the *DOG1* promoter (*DOGpro:LUC)* was tested by transient expression assays using agroinfiltration of tobacco (*Nicotiana benthamiana)* leaves (Figure 1E). Relatively strong luminescence signal, indicating substantial luciferase activity, and elevated levels of *LUC* mRNA were detected in leaves agroinfiltrated with *DOGpro:LUC* construct alone. However, luminescence and *LUC* mRNA expression from *DOGpro:LUC* was significantly reduced when co-infiltrated with an HSI2-expressing effector construct, indicating that expression from the *DOG1* promoter in tobacco leaves is repressed by co-expression of HSI2.

### B3 domain is required for HSI2 function

The B3 domain of HSI2 (B3_HSI2_) was reported to play an essential role in the regulation of *AGL15* and *FLC* by binding to RY elements in the proximal promoter or first intron of these genes, repectively (Qüesta et al., 2016; Yuan et al., 2016; Chen et al., 2018). We examined the function of the B3_HSI2_ domain in the regulation of the *DOG1* promoter by transient expression assays. An effector construct that encodes HSI2 with a mutated B3 domain (HSI2mB3) was coinfiltrated with the *DOG1pro:LUC* reporter construct into *N. benthamiana* leaves (Figure 2A). The *DOG1pro:LUC* reporter gene was strongly repressed by co-expression with intact HSI2; however, co-expression of HSI2mB3 resulted in high levels of reporter gene expression indicating that the B3 domain is required for HSI2-dependent repression of the *DOG1* promoter.

**Figure 2.**
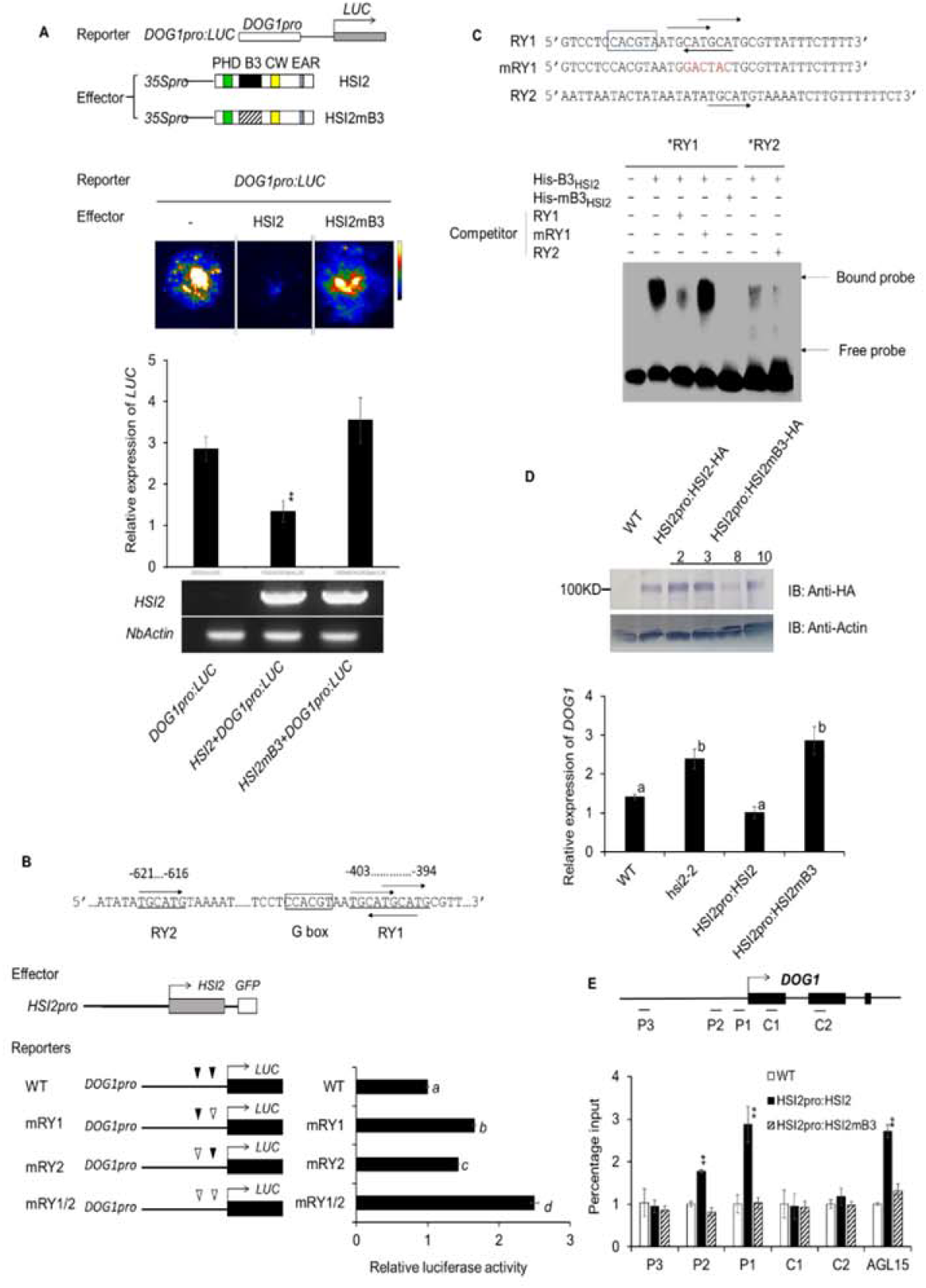
B3 domain is required for HSI2-dependent repression of *DOG1*. **(A)** Schematic representation of reporter and effector constructs used to assay the function of the HSI2 B3 domain. In the reporter construct, a luciferase gene is driven by the *DOG1* promoter. In effector constructs, expression of the intact HSI2 or HSI2mB3 is controlled by the *CaMV 35S* promoter, diagonal shading represents a mutated B3 domain. Luminescence images and relative expression of the luciferase mRNA from *N. benthamiana* leaves co-infiltrated with combinations of reporters and effectors, as indicated. *HSI2* and *NbActin* gene expression in infiltrated areas was assayed by RT-qPCR. **(B)** Function of RY1 and RY2 elements in the repression of the *DOG1* promoter. Schematic representation of the locations of RY1, RY2 and G-box sequence elements in the *DOG1* promotor are shown along with results from dual luciferase assays to evaluate the expression of reporter genes with intact or disrupted RY elements co-expressed with HSI2 in Arabidopsis protoplasts. **(C)** RY1, RY2, and mutant, mRY1 probes, shown with RY elements (arrows) and putative G-box (boxed) indicated, used in EMSA assays with His-B3HSI2 and mutant His-mB3_HSI2_ polypeptides. Bands representing free and bound probe are indicated. **(D)** Expression of HA-tagged HSI2mB3 protein was detected in transgenic Arabidopsis plants by immunoblot assays using anti-HA antibody, anti-actin was used to detect actin as a loading standard. Relative expression of *DOG1* in WT, *hsi2-2, HSI2pro:HSI2-HA* and *HSI2pro:HSI2mB3* Arabidopsis seedlings. RT-qPCR assays were normalized by *EF1a*. Lowercase letters indicate significant differences (*P*< 0.01) between genotypes. **(E)** Enrichment of HSI2-HA and HSI2mB3-HA at the *DOG1* locus in transgenic plants assayed by ChIP-qPCR. Tested regions are indicated in the gene structure. Assay for enrichment at the *AGL15* locus was included as a positive control. Data represent means of three ChIP-qPCR assays from three independent ChIP assays for each genotype. Error bars indicate SD. Asterisks indicate means significantly different from WT and *HSI2pro:HSI2mB3* at *P*< 0.01 (Student’s *t* test).

Sequence analysis indicated that RY elements are distributed in the *DOG1* proximal promoter between −621 and −394 bp upstream of the transcriptional start site. The region corresponding to P1 in Figure 2B contains overlapping RY elements (RY1,^5’^ TGCATGCATG^3’^), while the P2 region contains a single RY element (RY2,^5’^ TGCATG^3’^) (Figure 2B). The expression of luciferase reporter genes under control of the intact *DOG1* promoter or promoters with disruptions in RY1, RY2 or both were tested in Arabidopsis protoplasts using a dual luciferase assay. As show in Figure 2B, loss of both RY1 and RY2 resulted in a 2.5-fold increase in expression relative to the intact promoter, while mutation of RY1 led to a 2-fold increase and RY2 disruption gave a 1.5 fold increase. Thus, the *DOG1* RY elements appear to act additively with the RY1 element showing a somewhat stronger repressive effect than RY2.

To test whether the B3_HSI2_ domain can bind directly to the RY elements in the *DOG1* promoter *in vitro*, we performed electrophoretic mobility shift assay (EMSA). RY1- and RY2-containing DNA probes from the P1 and P2 regions of the *DOG1* promoter were labeled with biotin at the 3’ end and incubated with His-Trx-tagged B3_HSI2_ fusion protein. A strong signal indicating binding of B3_HSI2_ to probes containing the complex RY1 sequence element was detected, but the signal for binding to probe containing the RY2 element was much weaker (Figure 2C). Probe with a mutated RY1 element failed to bind B3_HSI2_, and a B3_HSI2_ polypeptide with a disrupted B3 domain (mB3_HSI2_) failed to bind the intact RY1 element-containing probe. These results indicate that the B3 domain of HSI2 can specifically bind, *in vitro*, to the complex RY1 element in the proximal promoter of *DOG1*, while binding to the single RY element found in the P2 region was substantially lower, relative to RY1, under the conditions used.

To test whether the B3 domain is required for HSI2 enrichment at the *DOG1* locus, we carried out ChIP-qPCR assays using a transgenic Arabidopsis line that expressed HSI2 with a disrupted B3 domain under the control of the native *HSI2* promoter in the *hsi2-2* mutant background (*HSI2pro:HSI2mB3)* (Chen et al., 2018). Plants from three independent transgenic lines (2, 3 and 10), that express HSI2mB3 at levels similar to the intact *HSI2* transgene were analyzed as biological replicates (Figure 2D). Expression of *DOG1* remained high in these lines, indicating that HSI2-dependent repression of *DOG1* was not restored by complementation with HSI2mB3. ChIP-qPCR analysis showed that, unlike intact HSI2, HSI2mB3 fails to accumulate at the proximal promoter region of the *DOG1* locus (Figure 2E). Similar results were obtained for the enrichment of HSI2 at the *AGL15* locus, which was included as a positive control in this ChIP-qPCR analysis.

### PHD domain is required for HSI2-mediated repression of *DOG1*

The HSI2 PHD domain (PHD_HSI2_) was shown to be essential for HSI2-dependent repression of *AGL15* expression and HSI2 enrichment at the *AGL15* locus (Chen et al., 2018). To test whether PHD_HSI2_ also plays a role in *DOG1* silencing, the *DOG1pro:LUC* reporter construct was coinfiltrated into *N. benthamiana* leaves with intact HSI2 or HSI2mPHD in which eight conserved amino acids of the PHD domain were substituted (C39S, C42S, C65S, C68S, H73L, C76S, C93S and C96S) (Chen et al., 2018). As expected, luciferase activity and mRNA expression levels were significantly reduced when *DOG1pro:LUC* was co-expressed with intact HSI2, but coexpression with HSI2mPHD had no effect on reporter gene expression (Figure 3A). Transgenic complementation of *hsi2-2* plants with HSI2mPHD also failed to repress *DOG1* expression (Figure 3B). Rather, expression of HSI2mPHD in *hsi2-2* plants resulted in an increase in *DOG1* expression by more than 2-fold, relative to the *hsi2-2* null allele, though expression in these plants was not as high as in *hsi2 hsl1* double mutant seedlings (Figure 3B). To test if the PHD domain is also required for HSI2 enrichment at the *DOG1* locus, we performed ChIP-qPCR using WT, and *HSI2pro:HSI2*- and *HSI2pro:HSI2mPHD*-expressing Arabidopsis seedlings (Figure 3C). The results showed that, relative to intact HSI2, HSI2mPHD failed to accumulate at the P1 or P2 regions of the *DOG1* gene, indicating that PHD domain is essential for HSI2 accumulation at the *DOG1* locus.

**Figure 3.**
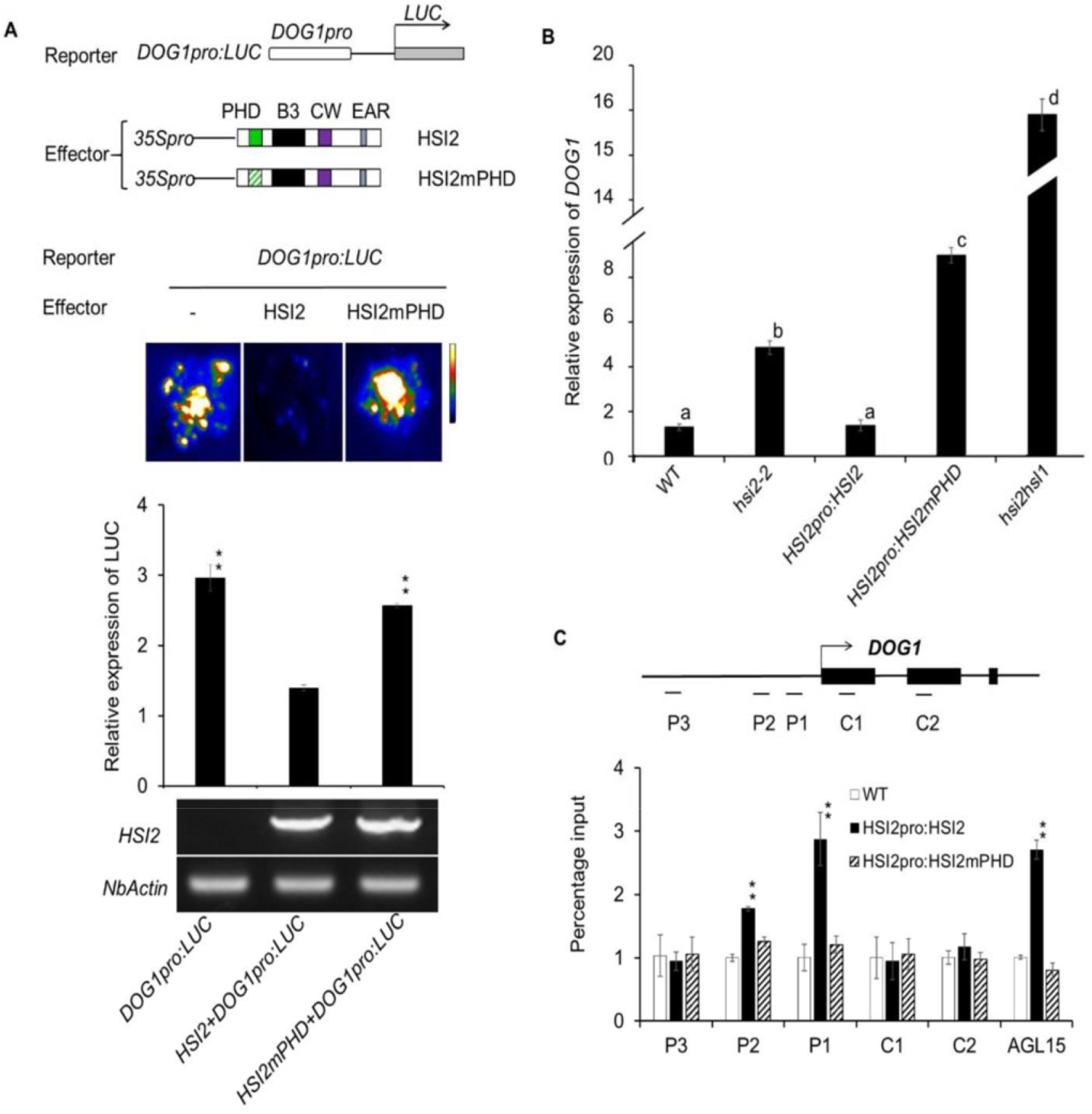
HSI2 PHD domain is essential for the repression of *DOG1*. **(A)** Schematic representation of reporter and effector constructs used to test the function of the HSI2 PHD domain. In effector constructs, green diagonal shading represents a mutated PHD domain. Luminescence image and relative expression of luciferase reporter genes from *N. benthamiana* leaves co-infiltrated with combinations of reporter and effector constructs, as indicated. *HSI2* and *NbActin* gene expression in each infiltrated area was assayed by RT-qPCR. **(B)** Relative expression of *DOG1* in Arabidopsis seedlings of WT, *hsi2-2, HSI2pro:HSI2-HA* and *HSI2pro:HSI2mPHD*. RT-qPCR assays were normalized using *EF1a*. Lowercase letters indicate significant differences (*P* < 0.01) between genetic backgrounds. **(C)** Results of ChlP-qPCR analyses to compare the enrichment of HSI2-HA and HSI2mPHD-HA at the *DOG1* locus in Arabidopsis plants. Tested regions are indicated in the gene structure. *AGL15* assay was included as a positive control. Data represent means of three ChlP-qPCR assays from three biological replicates. Error bars indicate SD. Asterisks indicate means significantly different from control plants (WT) at *P* < 0.01.

### CW and EAR domains are not required for HSI2-dependent repression of *DOG1*

Chen et al. (2018) reported that HSI2-mediated repression of *AGL15* expression requires EAR motif but not the CW domain. To investigate whether CW and EAR domains of HSI2 contribute to the repression of *DOG1*, we tested the functions of HSI2 with disrupted CW or deleted EAR domains in the regulation of the *DOG1* promoter by transient expression in *N. benthamiana* leaves (Supplementary Figure 1A). Similar to intact HSI2, co-expression of HSI2mCW or HSI2-ΔEAR effector constructs strongly repressed luciferase activity and decreased mRNA levels from the *DOG1pro:LUC* reporter construct, indicating that the CW and EAR domains are not required for HSI2-dependent repression of the *DOG1* promoter in agroinfiltrated leaves (Supplementary Figure 1B). Relative expression of endogenous *DOG1* was also tested in *HSI2pro:HSI2mCW* and *HSI2pro:HSI2mEAR* rescued *hsi2-2* Arabidopsis seedlings^27^. Consistent with the agroinfiltration assays, the results showed that the *DOG1* expression in these mutant lines was similar to that in WT seedlings, being strongly repressed relative to *hsi2-2* (Supplementary Figure 1C), confirming that intact HSI2 CW and EAR domains are not required for HSI2-mediated repression of *DOG1* expression in Arabidopsis.

### HSI2 and HSL1 form dimers *in vivo*

Chhun et al. (2016) reported the formation of HSI2:HSI2 and HSL1:HSL1 homodimers, along with HSI2:HSL1 heterodimers. To confirm these findings, we tested the interactions between HSI2 and HSL1 using both bimolecular fluorescence complementation (BiFC) and yeast two-hybrid assays, which showed both heterodimer and homodimer formation (Supplementary Figure 2A-C and Figure 4G). Further analysis indicated that the B3 domain is not required for dimerization since HSI2mB3 was still able to form homodimers (HSI2mB3:HSI2mB3), and heterodimer (HSI2mB3:HSL1) *in vivo* (Figure 4A, B). However, HSI2mPHD was unable to interact with either HSI2 or HSL1 (Figure 4C, D and G). These BiFC results were confirmed using both co-immunoprecipitation (CoIP) (Figure 4E, F) and yeast 2-hybrid assays (Figure 4G).

**Figure 4.**
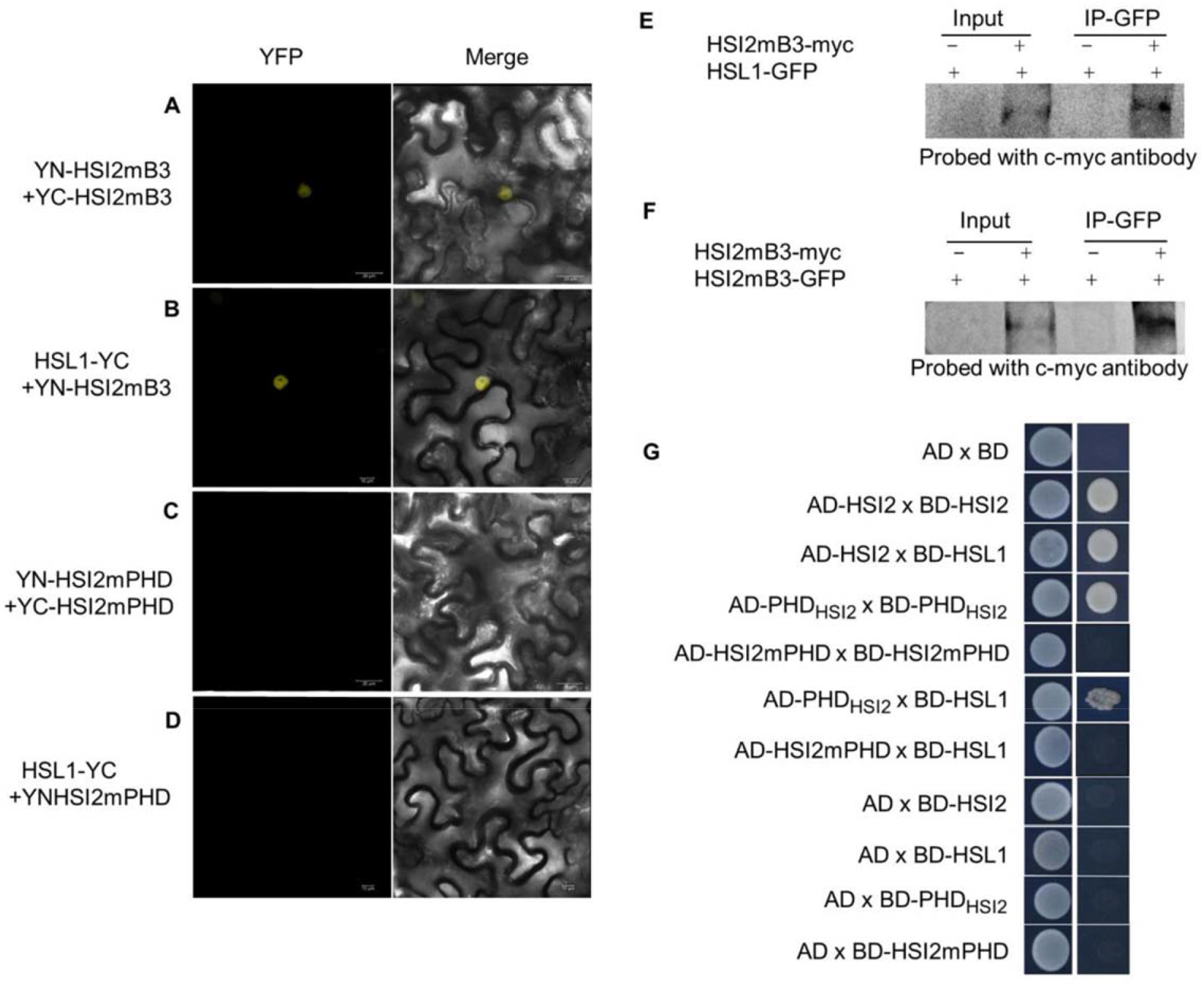
Dimerization between HSI2 and HSL1 is dependent on PHD-like domain. Bimolecular fluorescence complementation (BiFC) analysis of protein-protein interactions between **(A)** HSI2mB3:HSI2mB3, **(B)** HSI2mB3:HSL1, **(C)** HSI2mPHD:HSI2mPHD, and **(D)** HSI2mPHD:HSL1. **(E)** Co-immunoprecipitation (Co-IP) showing interaction between HSI2mB3 and HSI2mB3 and **(F)** HSI2mB3 and HSL1. Total protein was extracted from agroinfiltrated tobacco leaves co-expressing *35S:HSI2mB3-GFP* and *35S:HSI2mB3-myc, 35S:HSL1-GFP* and *35S: HSI2mB3-myc*, only *35S:HSI2mB3-GFP* or only *35S:HSL1-GFP*. HSI2mB3-GFP and HSL1-GFP was immunoprecipitated with anti-GFP antibody, and immunoblots were probed with anti-c-myc antibody. **(G)** Yeast two-hybrid interaction assays. Yeast cells transformed with indicated genes were selected on DDO (lacking Leu and Trp) and QDO (lacking adenine, His, Leu and Trp) media.

### HSL1 enrichment at the *DOG1* promoter is independent of HSI2

Protein sequence analysis showed that HSI2 and HSL1 share high sequence identity and contain similar conserved regions (Tsukagoshi et al., 2007). Since HSI2 harbors both PHD and B3 domains, which are required for HSI2 accumulation at the *DOG1* promoter (Figure 2E and 3C) and are essential for full repression of *DOG1* expression, we predicted that HSL1 might also be enriched in the chromatin of the *DOG1* locus. To confirm this, a gene construct that expresses GFP-tagged HSL1 under control of the native *HSL1* promoter (*HSL1pro:HSL1-GFP)* was transformed into *hsl1-1* Arabidopsis protoplast for ChIP-qPCR assays. The results indicated that, like HSI2, HSL1 is significantly enriched at the proximal promoter regions (P1 and P2), of the *DOG1* locus (Figure 5A). To test whether the enrichment of HSL1 at the *DOG1* locus depends on HSI2, ChIP-qPCR was also performed in protoplasts from *hsi2-2* Arabidopsis plants that transiently expressed HSL1-GFP. In these assays, HSL1 was still able to accumulate at the proximal promoter region of *DOG1* in the absence of HSI2, indicating that HSL1 enrichment at *DOG1* is independent of HSI2 (Figure 5B). Significant HSI2-independent enrichment of HSL1 was also detected at the *AGL15* promoter (Figure 5A, B). To confirm the hypothesis that HSL1 can physically bind to RY elements in the *DOG1* promoter *in vitro*, we performed EMSA assays using His-tagged HSL1-B3 fusion protein incubated with biotin-labelled probes containing RY1 and RY2 from the P1 and P2 regions of the *DOG1* promoter, respectively (Figure 5C). Consistent with the binding activity of the B3_HSI2_ polypeptide, the B3_HSL1_ polypeptide can bind probe that contains the complex RY1 element more effectively than to probe containing the single RY2 element (Figure 5C). However, EMSA signals for B3_HSL1_ binding to RY1 in these assays was much weaker than for B3_HSI2_. These results demonstrate that the B3 domain of HSL1 can bind to RY elements in the proximal promoter region of *DOG1*, leading to HSI2-independent HSL1 enrichment at the *DOG1* locus. However, the reduced EMSA signal for binding of B3_HSL1_ appears to indicate that its affinity for RY elements is reduced relative to the B3HSI2.

**Figure 5.**
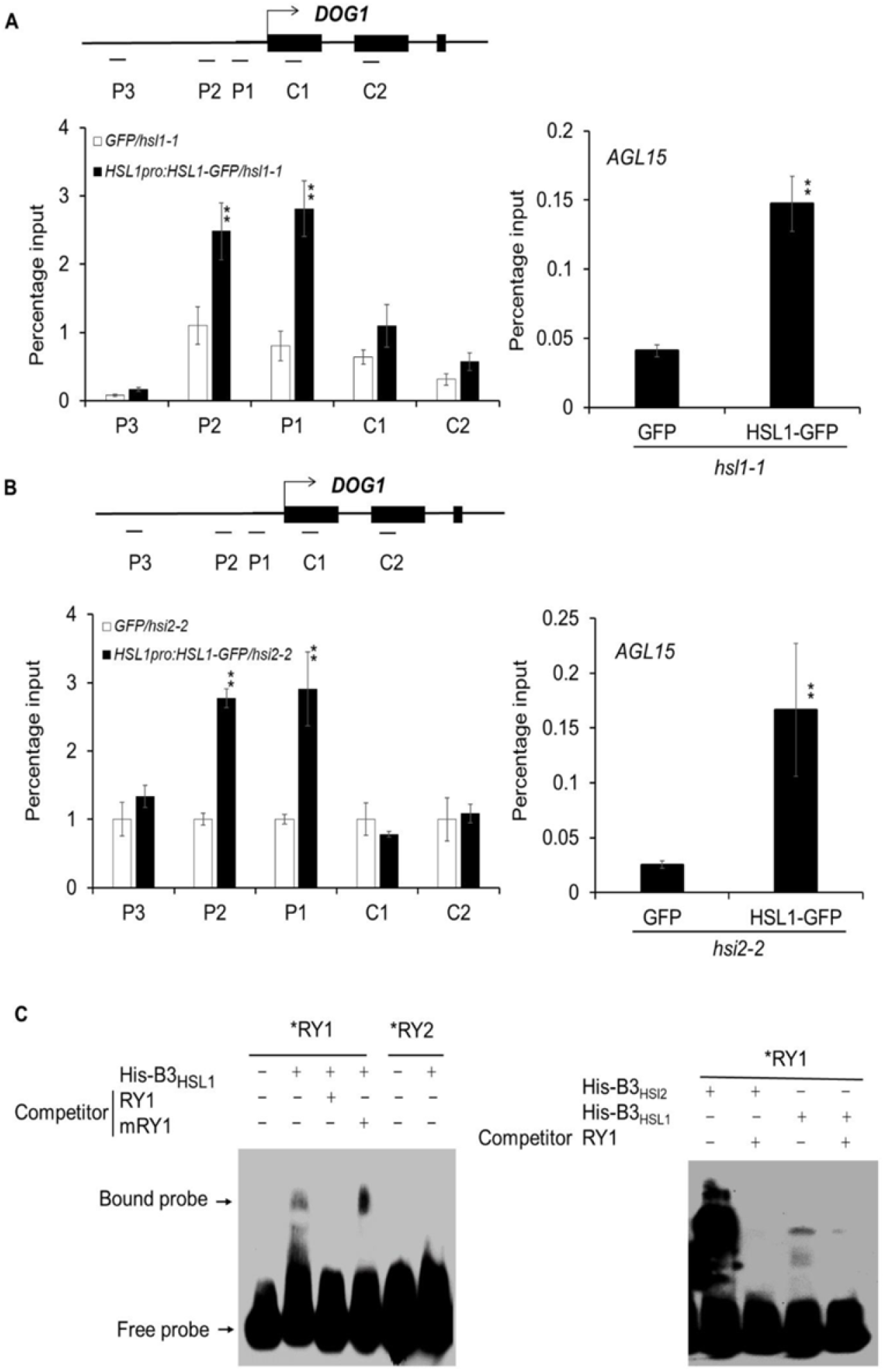
HSL1 directly binds to the *DOG1* promoter. **(A-B)** ChlP-qPCR analysis of HSL1 enrichment at *DOG1* locus in *hsl1-1* and *hsi2-2* Arabidopsis protoplasts. *HSL1-GFP* was expressed under control of the native *HSL1* promoter. Tested regions are indicated in the gene structure. ChlP-qPCR assays showing enrichment of HSL1 at the *AGL15* locus in both *hsl1-1* and *hsi2-2* genetic backgrounds are included as positive controls. Data represent means of three assays from three biological replicates. Error bars indicate SD. Asterisks indicate means significantly different from control at *P*<0.01. **(C)** EMSA assay for the detection of DNA-protein interaction between RY1 and RY2, and mutant, mRY1 probes with His-B3_HSL1_ polypeptide. Bands representing free and bound probe are indicated. The stronger EMSA signal seen when using the B3_HSI2_ polypeptide than with the B3_HSL1_ polypeptide indicates that the B3_HSI2_ domain may have higher affinity for the RY1 probe than the B3_HSL1_ domain.

### Repression of *DOG1* by HSI2 and HSL1 is associated with H3K27me3 enrichment

Previously, we showed that loss of HSI2 in *hsi2-2* knockout and disruption in *hsi2-4* PHD domain Arabidopsis mutant seedlings resulted in de-repression of *DOG1* expression and reduced deposition of H3K27me3 at the *DOG1* locus (Veerappan et al., 2012, 2014). Since HSL1 also targets *DOG1*, we used ChIP-qPCR to examine the effects of various *hsi2* and *hsl1* Arabidopsis mutations on the levels of H3K27me3 at *DOG1* (Figure 6). While enrichment of H3K27me3 was significantly reduced, relative to WT, across the *DOG1* locus in three *hsi2* mutant lines (*hsi2-2, HSI2mPHD* and *HSI2mB3)*, H3K27me3 levels in *hsl1-1* seedlings were similar to WT. These results indicate that intact HSI2 mediates the trimethylation of H3K27 at WT levels in the absence of HSL1 and the PHD and B3 domains are required for this activity. Thus, as with transcriptional repression, HSI2 is able to fully complement the loss of HSL1 in H3K27me3 deposition. However, H3K27me3 deposition at the *DOG1* locus was more strongly reduced in the *hsi2 hsl1* double mutant, than in the three *hsi2* mutants, with the greatest effect seen at the C1 region (Figure 6). Thus, it is apparent that HSL1 contributes to the deposition of H3K27me3 at this locus.

**Figure 6.**
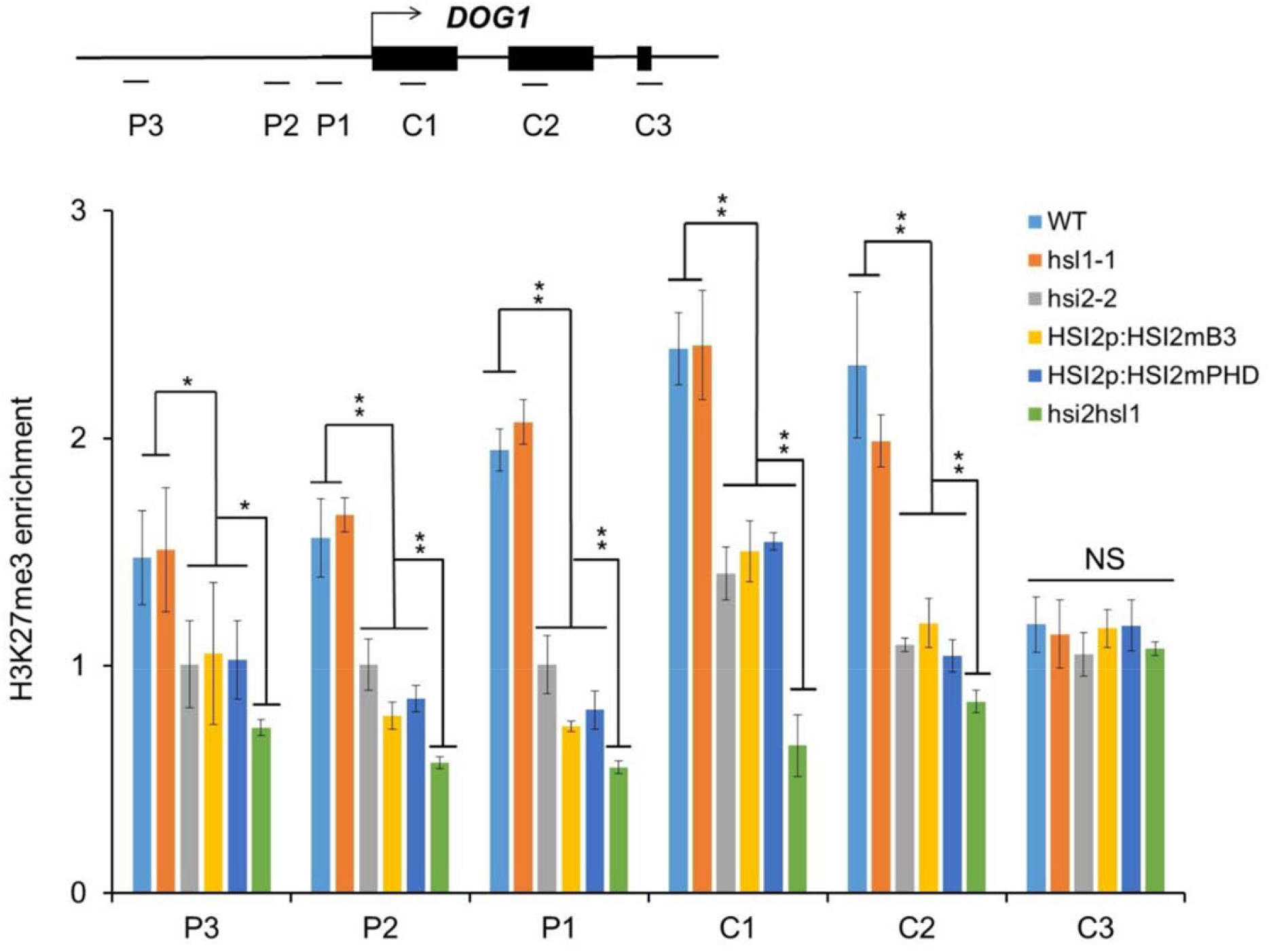
Both HSI2 and HSL1 are involved in deposition of H3K27me3 marks at *DOG1* locus. ChIP-qPCR analysis of H3K27me3 enrichment at the *DOG1* locus in WT, *hsl1-1, hsi2-2, HSI2pro:HSI2mPHD, HSI2pro:HSI2mB3, hsi2 hsl1* Arabidopsis seedlings. Tested regions are indicated in the gene structure. Data represent means of three assays from three biological replicates. Error bars indicate SD. * = *P*<0.05; ** = *P*<0.01; NS (non-significant).

### LHP1 and CLF interact with both HSI2 and HSL1 to regulate *DOG1* expression through deposition of H3K27me3

Yuan et al. (2016) reported that LHP1, which may serve as a bridge between PRC2 and PRC1 (Xu et al., 2008; Bratzel et al., 2010), is recruited by HSI2 to the *FLC* locus to promote deposition of H3K27me3, leading to downregulation of *FLC* expression. Therefore, it seemed likely that LHP1 could also be involved in the down-regulation of *DOG1* expression. To test this possibility, we evaluated the role of LHP1 in the repression of *DOG1*. Our results show that *DOG1* expression is significantly upregulated in *lhp1* Arabidopsis seeds, relative to that in WT, *hsl1-1* and *hsi2-2*, but remained lower than in the *hsi2 hsl1* double mutant (Figure 7A). Proteinprotein interactions between HSI2:LHP1 and HSL1:LHP1 *in vivo* were confirmed by BiFC and CoIP assays (Figure 7B, C) and BiFC analysis also showed that the B3_HSI2_ and B3_HSL1_ domains are sufficient for interaction with LHP1 (Figure 7D). ChIP-qPCR analysis using transgenic Arabidopsis seedlings that express Myc-LHP1 showed significant LHP1 enrichment at both the proximal promoter (P1) and exons (C1 and C2) of the *DOG1* locus (Figure 7E). Finally, ChIP-qPCR analysis in *lhp1* mutant seedlings showed significant reductions in H3K27me3 chromatin marks across the *DOG1* locus, relative to WT (Figure 7F). Shu et al. (2019) reported that CLF, a histone methyltransferase component of PRC2, accumulated at *FLC, AGL15* and *DOG1* genes that are directly regulated by HSI2 (Qüesta et al., 2016; Yuan et al., 2016; Chen et al., 2018). We used BiFC and CoIP assays to confirm that CLF interacts with HSI2 and HSL1 *in vivo* (Figure 7G) and ChIP-qPCR analysis of H3K27me3 deposition in *clf28* mutant seedlings showed significant decreases of this histone mark at P1, C1 and C2 regions of the *DOG1* locus compared to WT (Figure 7H). The relatively small effect on H3K27me3 in *clf28* could be due to the redundancy of CLF with the other PRC2 histone methyltransferase subunits SWINGER and MEDEA.

**Figure 7.**
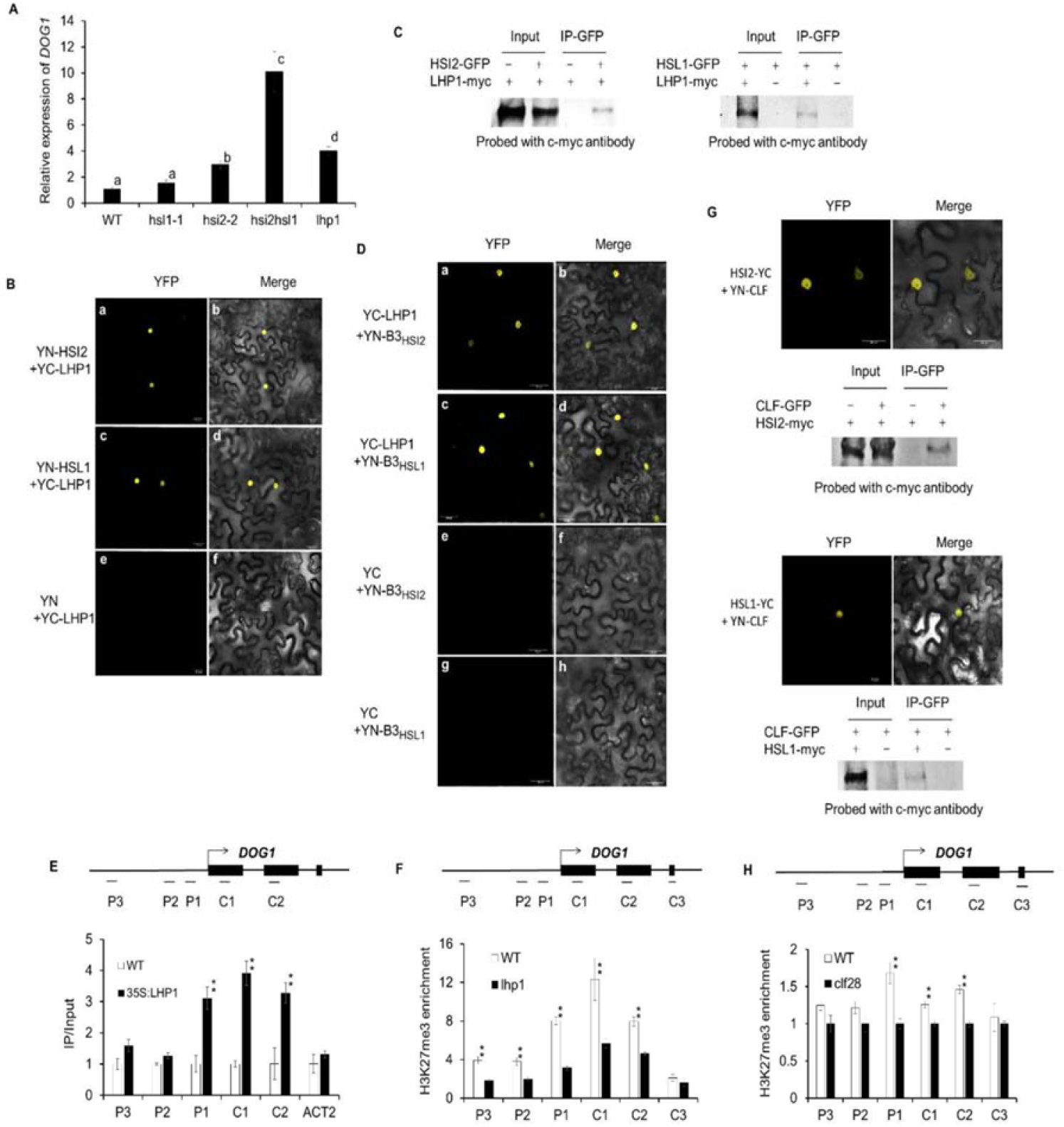
LHP1 and CLF interact with both HSI2 and HSL1. **(A)** Relative expression of *DOG1* in freshly-harvested seeds from WT, *hsl1-1, hsi2-2, hsi2 hsl1* Arabidopsis plants. RT-qPCR assays were normalized by *EF1a*. Lowercase letters indicate significant differences *P*<0.01. **(B)** BiFC analysis of LHP1:HSI2 and LHP1:HSL1 interactions. The N-terminal region of EYFP was fused to HSI2 (YN-HSI2) or HSL1 (YN-HSL1), and C-terminal region of EYFP was fused to LHP1 (YC-LHP1). EYFP fluorescence (YFP) and EYFP fluorescence images merged with bright-field images (Merge) are shown. Empty plasmid containing YN-YFP was used as a negative control (e, f). **(C)** Co-IP showing interaction of LHP1/HSI2, and LHP1/HSL1. Total protein was extracted from tobacco leaves co-expressing *35S:HSI2-GFP* and *35S:LHP1-myc, 35S:HSL1-GFP* and *35S:LHP1, 35S:HSL1-GFP* alone or *35S:LHP1-myc* alone. HSI2-GFP and HSL1-GFP was immunoprecipitated with anti-GFP antibody, and the immunoblot was probed with anti-c-myc antibody. **(D)** BiFC analysis showing that LHP1 interacts with the B3 domains of HSI2 (B3_HSI2_) and HSL1 (B3_HSL1_). **(E)** Results of ChIP-qPCR analysis of LHP1 enrichment at *DOG1* locus in WT and *35S:LHP1-myc* Arabidopsis seedlings. Tested regions are indicated in the gene structure. *ACT2* was included as a control. Data represent means of three assays from three biological replicates. Error bars indicate SD. Asterisks indicate means significantly different from control (WT) *P*<0.01. **(F)** Results of ChIP-qPCR analysis of H3K27me3 enrichment at *DOG1* locus in WT and *lhp1* Arabidopsis seedlings. **(G)** BiFC and Co-IP showing interaction of CLF/HSI2, and CLF/HSL. **(H)** Results of ChIP-qPCR analysis of H3K27me3 enrichment at *DOG1* locus in WT and *clf28* Arabidopsis seedlings.

## Discussion

We identified *DOG1*, a key regulator of seed dormancy in Arabidopsis (Alonso-Blanco et al., 2003; Bentsink et al., 2006, 2010; Huang et al., 2010), as a direct regulatory target of HSI2/HSL1-mediated transcriptional repression. HSI2 has been shown to regulate the developmental transition from seeds to seedlings, at least in part, by directly silencing the expression of the embryogenesis-promoting gene *AGL15* (Chen et al., 2018). HSI2 also affects the switch from vegetative growth to flowering by the down-regulation of *FLC* and *FLOWERING LOCUS T (FT)* (Qüesta et al., 2016; Yuan et al., 2016; Jing et al., 2019). In these cases, HSI2 was shown to interact, through its B3 DNA binding domain, with a pair of RY elements, located in the proximal promoters of *AGL15*, the first intron of *FLC* or the second intron of *FT*. We show here that HSI2 is also enriched in the chromatin at the proximal 5’ region of the *DOG1* locus and binds strongly, *in vitro*, to the complex RY1 region in the proximal *DOG1* promoter, which contains multiple overlapping canonical RY elements, and more weakly to the RY2 region, which contains a single RY element. (Figure 2B). Transient reporter gene expression analysis indicates that both RY1, and the simpler RY2 element contribute to transcriptional silencing and these observations agree with our ChIP-qPCR data that show strong enrichment of HSI2 and HSL1 *in vivo* at the P1 region, which contains the complex RY element, with lower but still significant enrichment at the P2 region, which contain the simple RY2 sequence. As with *AGL15* (Chen et al., 2018), the conserved HSI2 B3 domain is required for HSI2-dependent silencing of the *DOG1* promoter and enrichment of HSI2 at the *DOG1* locus *in vivo*.

Analysis of native *DOG1* expression in *hsi2* and *hsl1* Arabidopsis knock out mutant lines indicates that *HSI2* and *HSL1* act redundantly in the regulation of *DOG1* expression and seed dormancy (Figure 1A). While loss of HSL1 does not significantly effect *DOG1* expression, loss of HSI2 results in a 3-fold increase in expression of this gene. However, loss of both HSI2 and HSL1 leads to ~14-fold increase in *DOG1* expression and only the high levels of *DOG1* expression in the *hsi2 hsl1* double mutant resulted in a significant increase in seed dormancy. Therefore, the results presented here clearly indicate that both HSI2 and HSL1 are required for full transcriptional silencing of *DOG1* expression, which leads to the release of seed dormancy and early germination.

Our previously published data indicated that the PHD-like domain also plays a critical functional role in HSI2-mediated transcriptional silencing (Veerappan et al., 2012, 2014; Chen et al., 2018). As shown in Figure 2D, expression of HSI2mB3 in the *hsi2-2* seedlings fails to rescue the transcriptional repression of *DOG1*, resulting in expression levels similar to that in *hsi2-2* plants. On the other hand, expression of the HSI2mPHD mutant in the *hsi2-2* knock out mutant background results in *DOG1* expression reaching levels approximately 2-fold higher than in *hsi2-2* seedlings, though not as high as in the *hsi2 hsl1* double mutant (Figure 3B). This dominant-negative effect indicates that the PHD-like domain plays an important role in the functional interaction between HSI2 and HSL1. We confirmed that HSI2 and HSL1 can form homodimers and heterodimers *in vivo*, and the HSI2 PHD-like domain is required for dimerization. Therefore, it seems likely that the inability of HSI2mPHD subunits to dimerize interferes with HSL1 activity. One possible explanation for this could be competition between functionally intact HSL1 dimers and HSI2mPHD monomers, which have intact B3 domains and may retain the ability to bind non-productively to RY elements in the *DOG1* promoter. However, this seems unlikely since HSI2mPHD is not enriched at the *DOG1* locus (Figure 3C). Alternatively, HSI2mPHD monomers may compete with HSL1 dimers for components of the HSI2/HSL1 repressive complex such as PRC2 subunits, LHP1, and histone deacetylases.

*DOG1* expression is controlled by complex and diverse mechanisms involving alternative splicing, alternative polyadenylation, histone modifications, and a *cis*-acting antisense noncoding transcript known as *asDOG1* (Bentsink et al., 2006; Cyrek et al., 2016; Graeber et al., 2014; Müller et al., 2012; Fedak et al., 2016). Developmental regulation of *DOG1* expression during seed development depends on the LAFL network of master transcription factors that regulate seed maturation. These include a homolog of the NUCLEAR TRANSCRIPTION FACTORY subunit B known as LEAFY COTYLEDON1 (LEC1), along with ABSCISIC ACID INSENSITIVE3 (ABI3), FUSCA3 (FUS3), and LEC2, which, like HSI2, contain conserved, plant-specific B3 DNA binding domains that interact with RY *cis*-acting elements (Giraudat et al., 1992; Meinke et al., 1992; Keith et al., 1994; West et al., 1994; Lotan et al., 1998; Luerssen et al., 1998; Stone et al., 2001). Bryant et al. (2019) recently reported that increased expression of *DOG1* under low temperature conditions is directly regulated by bZIP67, and indirectly by LEC1.

These authors showed that bZIP67 binds to G box-like elements in the *DOG1* promoter, one of which is located just upstream of RY1 (Figure 2B) (Bryant et al., 2019). Disruption of this complex RY1 element in the *DOG1* promoter was reported to completely abolish its ability to drive *GUS* expression in protoplasts, suggesting that it is critical both for up-regulation and down-regulation of *DOG1* expression (Bryant et al., 2019). It is possible that the RY1 element could be bound by one or more of the LEC1-induced B3 domain-containing AFL factors ABI3, FUS3 or LEC2 (Stone et al., 2001; Braybrook et al., 2006; Baud et al., 2016) and is required for developmental regulation of *DOG1* expression (Braybrook et al., 2006; Baud et al., 2016; Pelletier et al., 2017; Mönke et al., 2012; Wang and Perry, 2013; González-Morales et al., 2016). This conclusion is supported by the results of ChIP experiments that showed that FUS3 binds to the *DOG1* locus *in vivo* (Wang and Perry, 2013). However, results from our transient expression assays using luciferase reporter genes controlled by intact and RY element-disrupted *DOG1* promoters showed that, rather than abolish promoter activity, loss of one or both RY elements resulted in increased reporter gene expression in protoplasts (Figure 3B). Therefore, the potential regulatory relationship between HSI2/HSL1 and AFL factors at the RY elements in the *DOG1* promoter remains to be elucidated.

Changes in chromatin structure associated with histone modifications play a critical role in the regulation of seed dormancy (Footitt et al., 2015). Repressive H3K27me3 marks form along the *DOG1* gene as dormancy declines and these marks accumulate rapidly on exposure to light. Our data support the hypothesis that HSI2 and HSL1 repress *DOG1* expression primarily through PRC2-dependent deposition of H3K27me3 at the *DOG1* locus. Compared to WT, three *hsi2* mutant lines, including the T-DNA knockout *hsi2-2*, and *hsi2-2* complemented with either *HSI2pro:HSI2mB3* or *HSI2pro:HSI2mPHD*, showed similar decreases in H3K27me3 levels at the *DOG1* locus. Although H3K27me3 at the *DOG1* locus in *hsl1* mutant seedlings was not significantly reduced relative to WT, H3K27me3 in *hsi2 hsl1* double mutant seedlings was substantially lower than in *hsi2* mutants. Therefore, although HSI2 is able to fully complement the *DOG1* expression and H3K27 trimethylation phenotypes in *hsl1* knockout seedlings, both HSI2 and HSL1 contribute to the deposition of H3K27me3 and transcriptional repression at the *DOG1* locus. Complementation of *hsi2-2* with *HSI2pro:HSI2mPHD* had a dominant-negative effect on the repression of *DOG1* expression (Figure 3B) but trimethylation of H3K27 in this line was not significantly different from *hsi2-2*. Therefore, it is possible that the HSI2 PHD domain is required to mediate, directly or indirectly, transcriptional repression mechanisms other than the deposition of H3K27me3. For example, the histone deacetylases HDA6 and HDA19 were reported to interact with HSI2 and HSL1, respectively (Chhun et al., 2016; Zhou et al., 2013), and Zeng et al. (2019) recently reported that HDA9 is required for polycomb-dependent silencing of *FLC*. These authors proposed that deacetylation of H3K27 by HDA9 may be necessary for its subsequent methylation by PRC2. Results reported by van Zanten et al. (2014) suggest that HDA9 negatively affects germination and is involved in the suppression of seedling traits in seeds, which is the opposite of that reported for HDA6 and HDA19, which repress embryonic characteristics in seedlings.

HSI2 directly interacts with the core PRC2 subunit MSI1 (Chen et al., 2018) and, as shown in Figure 7, both HSI2 and HSL1 also interact with CLF and LHP1. LHP1 interacts with PRC2 components, including MSI1, and is able to recognize H3K27me3 and spread this repressive histone mark throughout the Arabidopsis genome (Turck et al, 2007; Zhang et al, 2007; Exner et al., 2009; Derkacheva et al., 2013). HSI2 recruits LHP1 to the *FLC* locus (Yuan et al., 2016) and our results show that *DOG1* expression is de-repressed in *lhp1* seedlings (Figure 7A). LHP1 is enriched at the *DOG1* locus (Figure 7E), and is required for deposition of H3K27me3 chromatin marks (Figure 7F). Thus, it is clear that LHP1 plays an important role in the repression of *DOG1* and it is possible that LHP1 may provide a bridge between H3K27me3 marks and the HSI2/HSL1/PRC2 complex.

Based on our data, here we propose a model for the HSI2 and HSL1 mediated regulation of *DOG1* expression and seed dormancy shown in Figure 8. Stable repression of *DOG1* expression is mediated directly by the combined action of the closely related transcriptional repressors HSI2 and HSL1, which form dimers that bind, via their conserved B3 domains, to RY *cis*-acting elements located upstream of the transcription start site. Conserved PHD-like domains are necessary for dimer formation and are required for transcriptional silencing activity. Direct contacts with LHP1 and PRC2 subunits MSI1 and CLF lead to the formation of a repressive complex that represses *DOG1* expression through the accumulation and spread of H3K27me3 chromatin marks at the *DOG1* locus, which leads to the release of seed dormancy and promotion of germination. These data do not discount to potential functions of other transcriptional repressive cofactors such as histone deacetylases or MED13 in HSI2/HSL1-dependant silencing of *DOG1* and other direct target genes.

**Figure 8.**
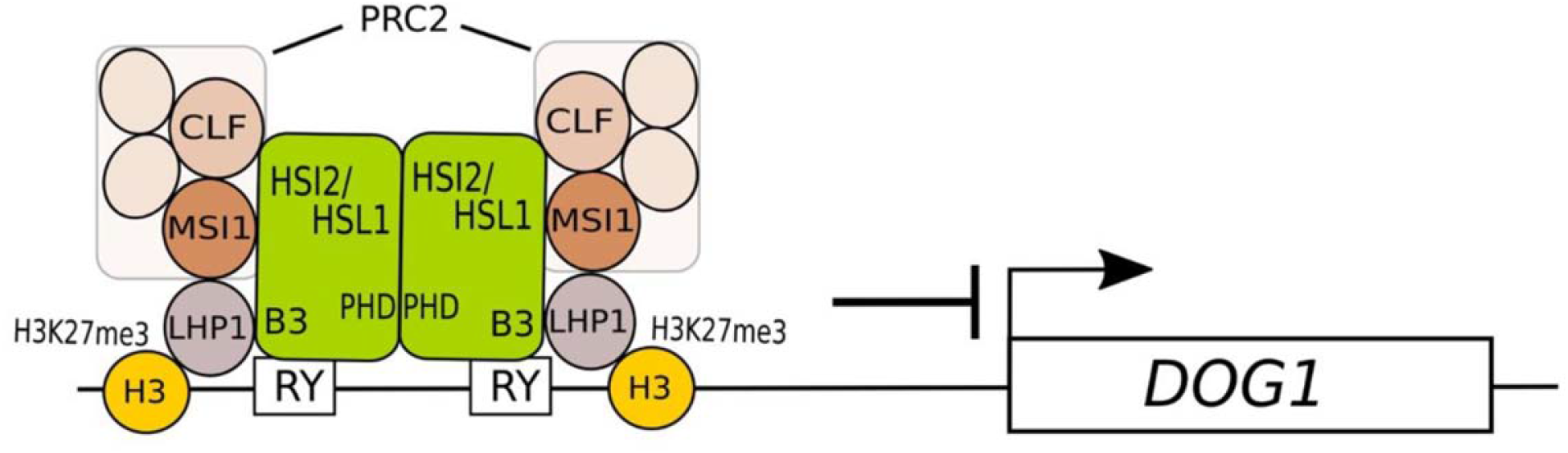
The proposed model for the repression of *DOG1* expression by HSI2 and HSL1. Simplified model showing the assembly of a hypothetical HSI2/HSL1 repressive complex at the *DOG1* locus. PHD domain-dependent HSI2/HSL1 dimerization allows for B3 domain-dependent binding to RY elements located upstream of the *DOG1* transcription start site (arrow). Direct interaction with LHP1 could provide links to H3K27me3 marks associated with nearby nucleosomes and PRC2 probably recruited through direct contacts with MSI1 and CLF subunits. The resulting complex, possibly along with other components, catalyzes the accumulation of additional H3K27me3 marks that repress *DOG1* expression, leading to the release of seed dormancy.

## Methods

### Plant Materials and Growth Conditions

Arabidopsis (*Arabidopsis thaliana)* Columbia (Col-0; CS60000) wild-type, loss-of-function alleles *hsl1-1* (SALK_059568), *lhp1-1* (CS3796) and *dog1-3* (SALK_000867), and gain-of-function allele *dog1-5* (SALK_022748) were obtained from Arabidopsis Biological Resources Center. For all the experiments, plants were grown under continuous illumination (fluorescent lamps at ~200μmol/m^2^s^2^) at 24°C on 0.3% Phytagel plates containing 0.5X Murashige and Skoog (MS) salt, 0.5 g/L MES, 1X Gamborg vitamin mix, and 1% sucrose (pH adjusted to 5.7).

### Plasmid Construct and Plant Transformation

The *HSL1* promoter, consisting of a 3687-bp fragment immediately upstream of the HSL1 start codon, was amplified by PCR and cloned into the binary vector pDONR207. After sequencing, the *HSL1* promoter was cleaved by ApaI restriction enzyme and inserted into pGWB504 containing *HSL1* cDNA to generate *HSL1pro:HSL1* DNA construct that contains a GFP tag at the C terminus. To generate *35Spro:LHP1* construct, pDONR207 containing the full-length *LHP1* coding sequence was subcloned into the destination vectors pGWB521 and pEarleygate104. Arabidopsis plants were transformed using the floral dip method (Clough and Bent, 1998). The primers used for generating these constructs are listed in Supplementary Data Set 1.

### Luciferase Imaging

Imaging of luciferase was performed using Andor iKON-M DU934N-BV CCD camera (Andor Technology). Andor SOLIS (I) imaging software (Andor Technology) was used for image acquisition and processing. Prior to luminescence imaging, Arabidopsis siliques and infiltrated *Nicotiana benthamiana* leaves were uniformly sprayed with 5mM D-luciferin (Gold Biotechnology) in 0.01% Triton X-100 solution. After spraying with luciferin, silique and leaf samples were incubated in the dark for 5 min. Exposure time for luminescence imaging was 10 min, unless otherwise specified.

### Transient Expression Assays

For reporter plasmids, a 1.5 Kb DNA sequence immediately upstream of the translational start codon of *DOG1* was cloned into pENTR/D TOPO vector, and then the *DOG1* promoter sequence and mutant derivatives of *DOG1* promoter (mRY1, mRY2 and mRY1/2) were subcloned into pGWB535 using the gateway system to generate reporter constructs. The construct that expresses HSI2pro:HSI2-GFP (Chen et al., 2018) was used as effector plasmid. Transient expression assays were performed with Arabidopsis protoplasts as described (Asai et al., 2002; Zhang et al., 2014). For each transformation, 5 μg of reporter and 4 μg of effector plasmid were used. For normalization of the activity of the reporter gene, 1 μg of plasmid pRLC was used as an internal control.

### RT-qPCR

RT-qPCR was performed using a StepOne Plus system (Applied Biosystems) with iTAqSYBR Green Supermix with ROX (Bio-Rad).RNase-free DNase (Qiagen)-treated total RNA was used for cDNA synthesis using iScript cDNA synthesis kit (Bio-Rad). For each experiment, cDNA synthesis reactions were performed on three independent RNA samples prepared from approximately five seedlings, each. Three qPCR reactions were performed for each cDNA sample. EF1A (AT5G60390) and HYGROMYCIN PHOSPHOTRANSFERASE (HPT) were used as reference genes for Arabidopsis and *N. benthamiana*, respectively. The relative expression of genes was calculated according to the ABI Prism 7700 sequence detection system (User Bulletin #2). Primer sequences used for RT-qPCR are listed in Supplementary Data Set 1.

### ChIP-qPCR Analysis

ChIP assays were performed as previously described^27^. Chromatin was extracted from seven day old seedlings grown in MS medium supplemented with 1% sucrose. The chromatin in these seedlings was cross-linked with 1% formaldehyde. The resulting chromatin was sheared to fragments with 500 bp (200–1000 bp) average length by sonication and used for immunoprecipitation with commercially available anti-GFP (Abcam, ab290), anti-HA (Abcam, ab9110) and anti-H3K27me3 (Millipore, 07-449), respectively. After reversing the cross-links, immunoprecipitated DNA was analyzed by qPCR using primers for specific regions of the *DOG1* gene. Three independent experiments, each using 500 mg of seedlings (25–30 individual plants), were performed. Three technical replications for each qPCR assay were performed, and *ACT2* was used as an internal control for normalization. In Arabidopsis leaf protoplasts, the ChIP assays were performed as described previously (Zhang et al., 2014; Lee et al., 2007; Du et al., 2009; Xiong et al., 2013). *HSL1pro:HSL1-GFP* DNA was transformed into *hsl1-1* and *hsi2-2* Arabidopsis protoplasts from 14 day old leaves using the polyethylene glycol-mediated transformation method. Protoplasts were incubated at room temperature for 13 h under dark conditions. Protoplast chromatin was cross-linked by 1% formaldehyde in W5 medium for 20 min and quenched with glycine (0.2 M) for 5 min. The protoplasts were then lysed, and the DNA was sheared on ice with sonication. Immunoprecipitation was performed with anti-GFP. After reversing the cross-links, the purified DNA was analyzed by qPCR using primers for specific regions of *DOG1*. Each experiment was repeated at least twice using protoplasts from approximately five leaves. Three technical replicates for each qPCR assay were performed, and *ACT2* was used as an internal control for normalization. Primers of target genes used for qPCR in ChIP analysis are listed in Supplementary Data Set 1.

### EMSA

EMSA was performed using double stranded biotinylated DNA probes and the Lightshift Chemiluminescent EMSA kit (Thermo Scientific). Biotin 3’ end-labeled DNA oligomers were prepared using a biotin end-labeling kit (Thermo Scientific), and double-stranded DNA probes were generated by annealing sense and antisense oligomers. Each 20-mL binding reaction contained 50 fmol of biotin-labeled double-stranded DNAs, 5 mg of recombinant His-HSI2B3 protein, and 50 ng/mL poly (dIdC) in binding buffer (10mM Tris, 30mM KCl, 0.1mM EDTA, 1mM DTT, 0.05% Nonidet P-40, and 6.5% glycerol, pH 7.9). Binding reactions were incubated for 20 min at room temperature resolved by electrophoresis of reaction samples on 5% polyacrylamide gels with TBE buffer. Detection of biotin-labeled DNA was performed by Andor iKON-M DU934N-BV CCD camera (Andor Technology).

### Yeast Two-Hybrid Assay

Yeast two-hybrid assays were performed using the Matchmaker GAL4-based two-hybrid system 3 (Clontech) according to the manufacturer’s instructions. Sequences that encode full-length HSI2, PHD_HSI2_ and HSI2mPHD were subcloned into the pGADT7 and pGBKT7 vectors, whereas the full-length HSL1 coding sequences were subcloned into the pGBKT7 vector. All constructs were transformed into yeast strain AH109 by the lithium acetate method, and yeast cells were grown on a minimal medium/-Leu/-Trp according to the manufacturer’s instructions (Clontech). Transformed colonies were plated onto a minimal medium/-Leu/-Trp/-His/-Ade to test for possible interactions.

### BiFC

Full-length cDNA of *LHP1* was cloned into pDONOR207 vector and subsequently introduced into pSITE-DEST-nEYFP-C1 vector containing the N-terminal fragment of YFP. The coding sequences of *HSL1, HSI2, HSI2mB3*, and *HSI2mPHD* were initially cloned into pENTR/SD/D-TOPO vector and subcloned into pSITE-DEST-nEYFP-C1 and pSITE-DEST-cEYFP-C1 vectors. *Agrobacterium tumefaciens* strain GV2260 carrying each construct was co-infiltrated into tobacco leaves. After two days, YFP fluorescence signal was observed by a laser scanning confocal microscope (Leica TCS SP8 confocal) at the Noble Research Institute Imaging Core Facility.

### *In Vivo* Co-immunoprecipitation (Co-IP) Assay

Total proteins from homogenized tobacco were extracted using extraction buffer (50 mM Tris-HCl, pH7.5, 150mMNaCl, 2mM EDTA, 10% glycerol, 0.2% 2-mercaptoethanol, 1% Triton-X 100, 1mM PMSF, and 1x protease inhibitor cocktail) and incubated 30 min at 4°C with gentle agitation. After centrifugation at 12,000 x g for 15min at 4°C, supernatant was incubated with protein A+G magnetic beads (Millipore 16-663) for 1h for preclearing at 4°C. The precleared protein solution was incubated with 50 μl Anti-GFP mAb-magnetic beads (MBL D153-11) overnight at 4°C. The supernatant was removed, and the magnetic beads were washed three times with extraction buffer. The proteins were eluted with 2X SDS sample loading buffer and analyzed by immunoblotting using anti-myc antibody (Thermo Fisher MA1-980).

## Author contribution

The research was performed and analyzed primarily by N.C. and he and R.D.A conceived the research, designed experiments and wrote the manuscript. Specific experiments were performed by H.W. and H.A. V.V. and M.T. provided critical analysis of the research and editing of the manuscript.

## Acknowledgments

We thank the Arabidopsis Biological Resource Center at Ohio State University for providing the T-DNA insertion mutants. We are also grateful to Dr. Jin Nakashima and Mihwa Yi at the Noble Research Institute for cellular fluorescence imaging. We also thank Dr. Kiran Mysore from the Noble Research Institute for critically reading the manuscript and for stimulating discussions. This work was supported by the Oklahoma Agricultural Experiment Station and Oklahoma Center for the Advancement of Science & Technology (Grant # PS16-009 to R.D.A).

---

**Supplementary Figure 1.** Disruption of CW and EAR domains does not affect HSI2-mediated regulation of *DOG1* expression.

**(A)** Schematic representation of reporter and effector used to test the function of the CW and EAR domains of HSI2. Effector constructs encodes either intact HSI2, HSI2mCW or HSI2-ΔEAR with an EAR motif deletion at C-terminus of HSI2. **(B)** Luminescence images and relative expression analysis of the *LUC* mRNA, by RT-qPCR, from *N. benthamiana* leaves coinfiltrated with combinations of reporter and effector constructs, as indicated. RT-PCR analysis of *HSI2* and *NbActin* gene expression in infiltrated areas as above. **(C)** Relative expression of *DOG1* in Arabidopsis seedlings of WT, *HSI2pro:HSI2-HA, HSI2pro:HSI2mCW* and *HSI2pro:mEAR*. RT-qPCR assays were normalized using *EF1A*. Lowercase letters indicate significant differences (P < 0.01).

**Supplementary Figure 2**. HSI2 and HSL1 form homodimers and heterodimers *in vivo*.

BiFC analysis of protein-protein interactions between **(A)** HSI2 and HSI2, **(B)**, HSI2 and HSL1 and **(C)** HSL1 and HSL1.

## Parsed Citations

Alonso-Blanco, C., Bentsink, L., Hanhart, C.J., Blankestijn-de Vries, H., and Koornneef, M. (2003). Analysis of natural allelic variation at seed dormancy loci of Arabidopsis thaliana. Genetics 164: 711–729.

Asai, T., Tena, G., Plotnikova, J., Willmann, M.R., Chiu, W.L., Gomez-Gomez, L., Boller, T., Ausubel, F.M., and Sheen, J. (2002). MAP kinase signaling cascade in Arabidopsis innate immunity. Nature 415: 977–983.

Baud, S., Kelemen, Z., Thévenin, J., Boulard, C., Blanchet, S., To, A., Payre, M., Berger, N., Effroy-Cuzzi, D., Franco-Zorrilla, J.M., Godoy, M., Solano, R., Thevenon, E., Parcy, F., Lepiniec, L., and Dubreucq, B. (2016) Deciphering the molecular mechanisms underpinning the transcriptional control of gene expression by master transcriptional regulators in Arabidopsis seed. Plant Physiol. 171: 1099–1112.

Bentsink, L., Jowett, J., Hanhart, C.J., and Koornneef, M. (2006). Cloning of DOG1, a quantitative trait locus controlling seed dormancy in Arabidopsis. Proceedings of the National Academy of Sciences, USA 103: 17042–17047.

Bentsink, L., Hanson, J., Hanhart, C.J., Blankestijn-de Vries, H., Coltrane, C., Keizer, P., El-Lithy, M., Alonso-Blanco, C., de Andrés, M.T., Reymond, M., van Eeuwijk, F., Smeekens, S., and Koornneef, M. (2010). Natural variation for seed dormancy in Arabidopsis is regulated by additive genetic and molecular pathways. Proc. Natl. Acad. Sci. USA 107(9):4264–9.

Bratzel, F., López-Torrejón, G., Koch, M., Del Pozo, J.C., and Calonje, M. (2010). Keeping cell identity in Arabidopsis requires PRC1 RING-finger homologs that catalyze H2A monoubiquitination. Curr. Biol. 20: 1853–1859.

Braybrook, S.A., Stone, S.L., Park, S., Bui, A.Q., Le, B.H., Fischer, R.L., Goldberg, R.B., and Harada, J.J. (2006). Genes directly regulated by LEAFY COTYLEDON2 provide insight into the control of embryo maturation and somatic embryogenesis. Proc. Natl. Acad. Sci. USA 103: 3468–3473.

Bryant, F.M., Hughes, D., Hassani-Pak, K., and Eastmond, P.J. (2019). Basic LEUCINE ZIPPER TRANSCRIPTION FACTOR67 transactivates DELAY OF GERMINATION1 to establish primary seed dormancy in Arabidopsis. Plant Cell 31: 1276–1288.

Chen, N., Veerappan, V., Abdelmageed, H., Kang, M., and Allen, R.D. (2018). HSI2/VAL1 Silences AGL15 to Regulate the Developmental Transition from Seed Maturation to Vegetative Growth in Arabidopsis. Plant Cell. 30: 600–619.

Chhun, T., Chong, S.Y., Park, B.S., Wong, E.C., Yin, J.L., Kim, M., and Chua, N.H. (2016). HSI2 repressor recruits MED13 and HDA6 to down-regulate seed maturation gene expression directly during Arabidopsis early seedling growth. Plant Cell Physiol. 57: 1689–706.

Chiang, G.C., Bartsch, M., Barua, D., Nakabayashi, K., Debieu, M., Kronholm, I., Koornneef, M., Soppe, W.J., Donohue, K., and De Meaux, J. (2011). DOG1 expression is predicted by the seed maturation environment and contributes to geographical variation in germination in Arabidopsis thaliana. Mol. Ecol. 20: 3336–3349.

Clough, S.J., and Bent, A.F. (1998). Floral dip: a simplified method for Agrobacterium-mediated transformation of Arabidopsis thaliana. Plant J. 16: 735–743.

Cyrek, M., Fedak, H., Cieslelski A., Guo, Y., Sliwa, A., Brzezniak, L., Krzyczmonik, K., Pietras, Z., Kaczanowski, S., Liu, F., and Swiezewski, S. (2016). Seed dormancy in Arabidopsis is controlled by alternative polyadenylation of DOG1. Plant Physiology 170, 947–955.

Derkacheva, M., Steinbach, Y., Wildhaber, T., Mozgová, I., Mahrez, W., Nanni, P., Bischof, S., Gruissem, W., and Hennig, L. (2013). Arabidopsis MSI1 connects LHP1 to PRC2 complexes. EMBO J. 32: 2073–2085.

Du, L., Ali, G.S., Simons, K.A., Hou, J., Yang, T., Reddy, AS., and Poovaiah, B.W. (2009). Ca2+/calmodulin regulates salicylic-acid mediated plant immunity. Nature 457: 1154–1158.

Exner, V., Aichinger, E., Shu, H., Wildhaber, T., Alfarano, P., Caflisch, A., Gruissem, W., Köhler, C. and Hennig, L. (2009). The chromodomain of LIKE HETEROCHROMATIN PROTEIN 1 is essential for H3K27me3 binding and function during Arabidopsis development. PLoS One 4: e5335.

Fedak, H., Palusinska, M., Krzyczmonik, K., Brzezniak, L., Yatusevich, R., Pietras, Z., Kaczanowski, S., and Swiezewski, S. (2016). Control of seed dormancy in Arabidopsis by a cis-acting noncoding antisense transcript. Proc. Natl. Acad. Sci. USA 113: E7846–E7855.

Finch-Savage, W.E., and Leubner-Metzger, G. (2006). Seed dormancy and the control of germination. New Phytologist 171: 501–523.

Footitt, S., Müller, K., Kermode, A.R., and Finch-Savage, W.E. (2015) Seed dormancy cycling in Arabidopsis: chromatin remodelling and regulation of DOG1 in response to seasonal environmental signals. Plant J. 81: 413–442.

Giraudat, J., Hauge, B.M., Valon, C., Smalle, J., Parcy, F., and Goodman, H.M. (1992). Isolation of the Arabidopsis ABI3 gene by positional cloning. Plant Cell 4: 1251–1261.

González-Morales, S.I., Chávez-Montes, R.A., Hayano-Kanashiro, C., Alejo-Jacuinde, G., Rico-Cambron, T.Y., de Folter, S., and Herrera-Estrella, L. (2016). Regulatory network analysis reveals novel regulators of seed desiccation tolerance in Arabidopsis thaliana. Proc. Natl. Acad. Sci. USA 113: E5232–E5241.

Graeber, K., Linkies, A., Steinbrecher, T., Mummenhoff, K., Tarkowská, D., Turečková, V., Ignatz, M., Sperber, K., Voegele, A., de Jong, H., Urbanová, T., Strnad, M., and Leubner-Metzger, G. (2014). DELAY OF GERMINATION 1 mediates a conserved coat-dormancy mechanism for the temperature-and gibberellin-dependent control of seed germination. Proceedings of the National Academy of Sciences, USA 111, E3571–3580.

Huang, X., Schmitt, J., Dorn, L., Griffith, C., Effgen, S., Takao, S., Koornneef, M., and Donohue, K. (2010). The earliest stages of adaptation in an experimental plant population: strong selection on QTLs for seed dormancy. Molecular Ecology 19: 1335–1351.

Huo, H., Wei, S., and Bradford, K.J. (2016). DELAY OF GERMINATION1 (DOG1) regulates both seed dormancy and flowering time through microRNA pathways. Proceedings of the National Academy of Sciences, USA 113, E2199–E2206.

Jing, Y., Guo, Q., and Lin, R. (2019). The B3-Domain Transcription Factor VAL1 Regulates the Floral Transition by Repressing FLOWERING LOCUS T. Plant Physiol. 181: 236–248.

Keith, K., Kraml, M., Dengler, N.G., and McCourt, P. (1994). fusca3: A heterochronic mutation affecting late embryo development in Arabidopsis. Plant Cell 6: 589–600.

Kendall, S.L., Hellwege, A., Marriot, P., Whalley, C., Graham, I.A., and Penfield, S. (2011). Induction of dormancy in Arabidopsis summer annuals requires parallel regulation of DOG1 and hormone metabolism by low temperature and CBF transcription factors. Plant Cell 23: 2568–2580.

Koornneef, M., Bentsink, L., and Hilhorst, H. (2002). Seed dormancy and germination. Current Opinion in Plant Biology 5: 33–36.

Lafos, M., Kroll, P., Hohenstatt, M.L., Thorpe, F.L., Clarenz, O., and Schubert, D. (2011). Dynamic regulation of H3K27 trimethylation during Arabidopsis differentiation. PLoS Genet. 7: e1002040.

Lee, J.H., Yoo, S.J., Park, S.H., Hwang, I., Lee, J.S., and Ahn, J.H. (2007). Role of SVP in the control of flowering time by ambient temperature in Arabidopsis. Genes Dev. 21: 397–402.

Li, X., Chen, T., Li, Y., Wang, Z., Cao, H., Chen, F., Li, Y., Soppe, W.J.J., Li, W., and Liu, Y. (2019). ETR1/RDO3 regulates seed dormancy by relieving the inhibitory effect of the ERF12-TPL complex on DELAY OF GERMINATION1 expression. Plant Cell: 832–847.

Libault, M., Tessadori, F., Germann, S., Snijder, B., Fransz, P., and Gaudin, V. (2005). The Arabidopsis LHP1 protein is a component of euchromatin. Planta 222: 910–925.

Liu, Y., Ye, N., Liu, R., Chen, M., and Zhang, J. (2010). H2O2 mediates the regulation of ABA catabolism and GA biosynthesis in Arabidopsis seed dormancy and germination. Journal of Experimental Botany 61: 2979–2990.

Lotan, T., Ohto, M., Yee, K.M., West, M.A., Lo, R., Kwong, R.W., Yamagishi, K., Fischer, R.L., Goldberg, R.B., and Harada, J.J. (1998). Arabidopsis LEAFY COTYLEDON1 is sufficient to induce embryo development in vegetative cells. Cell 93, 1195–1205.

Luerssen, H., Kirik, V., Herrmann, P., and Misera, S. (1998). FUSCA3 encodes a protein with a conserved VP1/AB13-like B3 domain which is of functional importance for the regulation of seed maturation in Arabidopsis thaliana. Plant J. 15: 755–764.

Meinke, D.W. (1992). A homeotic mutant of Arabidopsis thaliana with leafy cotyledons. Science 258: 1647–1650.

Molitor, A.M., Bu, Z., Yu, Y., and Shen, W.H. (2014). Arabidopsis AL PHD-PRC1 complexes promote seed germination through H3K4me3-to-H3K27me3 chromatin state switch in repression of seed developmental genes. PLoS Genet. 10: e1004091.

Mönke, G., Seifert, M., Keilwagen, J., Mohr, M., Grosse, I., Hähnel, U., Junker, A., Wsisshaar, B., Conrad, U., Bäumlein, H., and Altschmied, L. (2012). Toward the identification and regulation of the Arabidopsis thaliana ABI3 regulon. Nucleic Acids Res. 40: 8240–8254.

Müller, K., Bouyer, D., Schnittger, A., and Kermode, A.R. (2012). Evolutionarily conserved histone methylation dynamics during seed life-cycle transitions. PLoS One 7: e51532.

Nakabayashi, K., Bartsch, M., Xiang, Y., Miatton, E., Pellengahr, S., Yano, R., Seo, M., and Soppe, W.J. (2012). The time required for dormancy release in Arabidopsis is determined by DELAY OF GERMINATION1 protein levels in freshly harvested seeds. Plant Cell 24: 2826–2838.

Née, G., Kramer, K., Nakabayashi, K., Yuan, B., Xiang, Y., Miatton, E., Finkemeier, I., and Soppe, WJJ. (2017). DELAY OF GERMINATION1 requires PP2C phosphatases of the ABA signaling pathway to control seed dormancy. Nat. Commun. 13;8(1):72.

Pelletier, J.M., Kwong, R.W., Park, S., Le, B.H., Baden, R., Cagliari, A., Hashimoto, M., Munoz, M.D., Fischer, R.L., Goldberg, R.B., and Harada, J.J. (2017). LEC1 sequentially regulates the transcription of genes involved in diverse developmental processes during seed development. Proc. Natl. Acad. Sci. USA 114: E6710–E6719.

Qüesta, J.I., Song, J., Geraldo, N., An, H., and Dean, C. (2016). Arabidopsis transcriptional repressor VAL1 triggers polycomb silencing at FLC during vernalization. Science 353: 485–488.

Shu, J., Chen, C., Thapa, R.K., Bian, S., Nguyen, V., Yu, K., Yuan, Z.C., Liu, J., Kohalmi, S.E., Li, C., and Cui, Y. 2019. Genome-wide occupancy of histone H3K27 methyltransferases CURLY LEAF and SWINGER in Arabidopsis seedlings. Plant Direct 3: e00100. doi: 10.1002/pld3.100.

Stone, S.L., Kwong, L.W., Yee, K.M., Pelletier, J., Lepiniec, L., Fischer, R.L., Goldberg, R.B., and Harada, J.J. (2001). LEAFY COTYLEDON2 encodes a B3 domain transcription factor that induces embryo development. Proc. Natl. Acad. Sci. USA 98: 11806–11811.

Suzuki, M., Wang, H.H., and McCarty, D.R. (2007). Repression of the LEAFY COTYLEDON 1/B3 regulatory network in plant embryo development by VP1/ABSCISIC ACID INSENSITIVE 3-LIKE B3 genes. Plant Physiol. 143: 902–911.

Tsukagoshi, H., Saijo, T., Shibata, D., Morikami, A., and Nakamura, K. (2005). Analysis of a sugar response mutant of Arabidopsis identified a novexl B3 domain protein that functions as an active transcriptional repressor. Plant Physiol. 138: 675–685.

Tsukagoshi, H., Morikami, A., and Nakamura, K. (2007). Two B3 domain transcriptional repressors prevent sugar-inducible expression of seed maturation genes in Arabidopsis seedlings. Proc. Natl. Acad. Sci. USA 104: 2543–2547.

Turck, F., Roudier, F., Farrona, S., Martin-Magniette, M.L., Guillaume, E., Buisine N, Gagnot, S., Martienssen, R.A., Coupland, G., and Colot, V. (2007). Arabidopsis TFL2/LHP1 specifically associates with genes marked by trimethylation of histone H3 lysine 27. PLoS Genet. 3: e86.

van Zanten, M., Zöll, C., Wang, Z., Philipp, C., Carles, A., Li, Y., Kornet, N.G., Liu, Y., and Soppe, W.J. (2014). HISTONE DEACETYLASE 9 represses seedling traits in Arabidopsis thaliana dry seeds. Plant J. 80: 475–488.

Veerappan, V., Wang, J., Kang, M., Lee, J., Tang, Y., Jha, A.K., Shi, H., Palanivelu, R., and Allen, R.D. (2012). A novel HSI2 mutation in Arabidopsis affects the PHD-like domain and leads to de-repression of seed-specific gene expression. Planta 236: 1–17.

Veerappan, V., Chen, N., Reichert, A.I., and Allen, R.D. (2014). HSI2/VAL1 PHD-like domain promotes H3K27 trimethylation to repress the expression of seed maturation genes and complex transgenes in Arabidopsis seedlings. BMC Plant Biol. 14: 293. doi: 10.1186/s12870-014-0293-4.

Veluchamy, A., Jégu, T., Ariel, F., Latrasse, D., Mariappan, K.G., Kim, S.K., Crespi, M., Hirt, H., Bergounioux, C., Raynaud, C., and Benhamed, M. (2016). LHP1 Regulates H3K27me3 Spreading and Shapes the Three-Dimensional Conformation of the Arabidopsis Genome. PLoS One 11: e0158936.

Wing, F., and Perry, S.E. (2013). Identification of direct targets of FUSCA3, a key regulator of Arabidopsis seed development. Plant Physiol. 161: 1251–1264.

West, M.A.L., Matsuderai Yee, K., Danao, J., Zimmerman, J.L., Fisher, R.L., Goldberg, R.B., and Harada, J.J. (1994). LEAFY COTYLEDON1 is an essential regulator of late embryogenesis and cotyledon identity in Arabidopsis. Plant Cell 6: 1731–1745.

Xiong, Y., McCormack, M., Li, L., Hall, Q., Xiang, C., and Sheen, J. (2013). Glucose-TOR signaling reprograms the transcriptome and activates meristems. Nature 496: 181–186.

Xu, L., and Shen, W.-H. (2008). Polycomb silencing of KNOX genes confines shoot stem cell niches in Arabidopsis. Curr. Biol. 18: 1966–1971.

Yuan, W., Luo, X., Li, Z., Yang, W., Wang, Y., Liu, R., Du, J., and He, Y. (2016). A cis cold memory element and a trans epigenome reader mediate Polycomb silencing of FLC by vernalization in Arabidopsis. Nature Genetics 48: 1527–1534.

Zeng, X., Gao, Z., Jiang, C., Yang, Y., Lie, R., and He, Y. (2019). HISTONE DEACETYLASE 9 functions with Polycomb silencing to repress FLOWERING LOCUS C expression. Plant Physiol. DOI:10.1104/pp.19.00793

Zhang, X., Germann, S., Blus, B.J., Khorasanizadeh, S., Gaudin, V. and Jacobsen, S.E. (2007). The Arabidopsis LHP1 protein colocalizes with histone H3 Lys27 trimethylation. Nat Struct Mol Biol. 14: 869–71.

Zhang, F., Wang, Y., Li, G., Tang, Y., Kramer, E.M., and Tadege, M. (2014). STENOFOLIA recruits TOPLESS to repress ASYMMETRIC LEAVES2 at the leaf margin and promote leaf blade outgrowth in Medicago truncatula. Plant Cell 26: 650–664.

Zhou, Y., Tan, B., Luo, M., Li, Y., Liu, C., Chen, C., Yu, CW., Yang, S., Dong, S., Ruan, J., Yuan, L., Zhang, Z., Zhao, L., Li, C., Chen, H., Cui, Y., Wu, K., and Huang, S. (2013) HISTONE DEACETYLASE19 interacts with HSL1 and anticipates in the repression of seed maturation genes in Arabidopsis seedlings. Plant Cell 25: 134–148.

